# Yeast volatomes differentially effect larval feeding in an insect herbivore

**DOI:** 10.1101/721845

**Authors:** Joel Ljunggren, Felipe Borrero-Echeverry, Amrita Chakraborty, Tobias U. Lindblom, Erik Hedenström, Maria Karlsson, Peter Witzgall, Marie Bengtsson

## Abstract

Yeasts form mutualistic interactions with insects. Hallmarks of this interaction include provision of essential nutrients, while insects facilitate yeast dispersal and growth on plant substrates. A phylogenetically ancient, chemical dialogue coordinates this interaction, where the vocabulary, the volatile chemicals that mediate the insect response, remains largely unknown. Here, we employed gas chromatography-mass spectrometry (GC-MS), followed by hierarchical cluster (HCA) and orthogonal partial least square discriminant analysis (OPLS-DA), to profile the volatomes of six *Metschnikowia* spp., *Cryptococcus nemorosus* and brewer’s yeast *Saccharomyces cerevisiae*. The yeasts, which are all found in association with insects feeding on foliage or fruit, emit characteristic, species-specific volatile blends that reflect the phylogenetic context. Species-specificity of these volatome profiles aligned with differential feeding of cotton leafworm larvae *Spodoptera littoralis* on these yeasts. Bioactivity correlates with yeast ecology; phylloplane species elicited a stronger response than fruit yeasts, and larval discrimination may provide a mechanism for establishment of insect-yeast associations. The yeast volatomes contained a suite of insect attractants known from plant and especially floral headspace, including (*Z*)-hexenyl acetate, ethyl (2*E*,4*Z*)-deca-2,4-dienoate (pear ester), (3E)-4,8-dimethylnona-1,3,7-triene (DMNT), linalool, α-terpineol, β-myrcene or (*E,E*)-a-farnesene. A wide overlap of yeast and plant volatiles, notably floral scents further emphasizes the prominent role of yeasts in plant-microbe-insect relationships including pollination. The knowledge of insect-yeast interactions can be readily brought to practical application, live yeasts or yeast metabolites mediating insect attraction provide an ample toolbox for the development of sustainable insect management.

**IMPORTANCE:** Yeasts interface insect herbivores with their food plants. Communication depends on volatile metabolites, and decoding this chemical dialogue is key to understanding the ecology of insect-yeast interactions. This study explores the volatomes of eight yeast species which have been isolated from foliage, flowers or fruit, and from plant-feeding insects. They each release a rich bouquet of volatile metabolites, including a suite of known insect attractants from plant and floral scent. This overlap underlines the phylogenetic dimension of insect-yeast associations, which according to the fossil record, long predate the appearance of flowering plants. Volatome composition is characteristic for each species, aligns with yeast taxonomy, and is further reflected by a differential behavioural response of cotton leafworm larvae, which naturally feed on foliage of a wide spectrum of broad-leaved plants. Larval discrimination may establish and maintain associations with yeasts and is also a substrate for designing sustainable insect management techniques.

---

Microorganisms essentially interface plants and insects (1-4). Invisibly, microorganisms broadcast a rich bouquet of volatile metabolites to mediate communication with other microorganisms, plants and associated animals (5-7). Plants also release volatile compounds in abundance, but there is mounting evidence that the production of volatiles by bacteria (8, 9), fungi (10-12) and yeasts (13-17) is equally prolific. A wide overlap in compounds released by plants (18) and microbes (19) suggests furthermore that plant headspace includes volatiles that are produced by plant-associated epiphytic and endophytic microbes. A striking example is floral scent, where bacteria and especially yeasts metabolize pollen, nectar and other floral compounds and thus become a prominent source of volatiles (20-25).

Yeasts are widely associated with insects since they require vectors for dispersal and outbreeding. In addition, larval feeding facilitates yeast growth on plant substrate. Yeasts provide, on the other hand, nutritional services to insects (26-32). This mutualistic interaction is facilitated by communication with volatile metabolites. Yeasts have apparently evolved the capacity to synthesize aroma compounds to attract insects (33-35) and insects, on the other hand, possess dedicated olfactory receptors tuned to yeast fermentation metabolites that signal suitable substrates for adult and larval feeding (36-39). Consequently, yeast volatiles, in addition to plant volatiles, play a part in host-plant and food finding in insect herbivores, including flies and moths (5, 40-44).

Investigations of the volatile metabolites of insect-associated yeasts serve a dual purpose. Plants and their insect herbivores are fundamental to many ecosystems. Olfactory recognition of food plants by insects is key to their interactions (45-47), and it is, accordingly, of fundamental interest to identify the microbial component of plant-insect communication. Furthermore, yeast metabolites or live yeasts facilitate insect management. Lack of environmentally benign, yet efficient control methods is an increasingly pressing issue in times of global change and increasing food insecurity (48-52). Yeast attraction of insects and their larvae for feeding (33, 44, 53-56) can be exploited for population control of herbivores (57, 58), as well as for improved crop pollination (59).

We asked whether taxonomically related yeasts, isolated from different insects and habitats, differ with respect to their volatile metabolomes, and whether cotton leafworm larvae behave differently towards them. We selected a *Cryptococcus* and several *Metschnikowia* yeasts, which have been isolated from insect larvae feeding on fruit or foliage, including the cotton leafworm, *Spodoptera littoralis*, a polyphagous noctuid moth (60). We have investigated the volatomes of these yeasts by gas chromatography-mass spectrometry (GC-MS) and comparative multivariate discriminant analysis, affording unique volatile fingerprints. Differential larval attraction reveals the behavioural relevance of characteristic differences in yeast volatome composition.

## RESULTS

### Yeast headspace analysis

A phylogenetic tree of the yeasts investigated, *Cryptococcus nemorosus*, six *Metschnikowia* spp. and baker’s yeast *Saccharomyces cerevisiae* is shown in Figure 1. These yeasts have all been found to occur in association with insects. The volatiles released during fermentation were investigated by GC-MS, and shown to contain a range of compounds, including largely methyl and ethyl esters but also terpenoids, straight and branched alcohols, aldehydes, ketones, acids, five sulfur and three nitrogen-containing volatiles (Figure 2, Supplemental Table S1). Twenty-six of the 192 compounds found are not yet listed in databases of volatiles from yeasts, fungi and bacteria, and 33 compounds are new for yeasts (17, 19). Many of these yeast-produced compounds, including terpenoids such as linaool and farnesenes, and esters, such as pear ester, have also been found in plant headspace (18).

**FIG 1.**
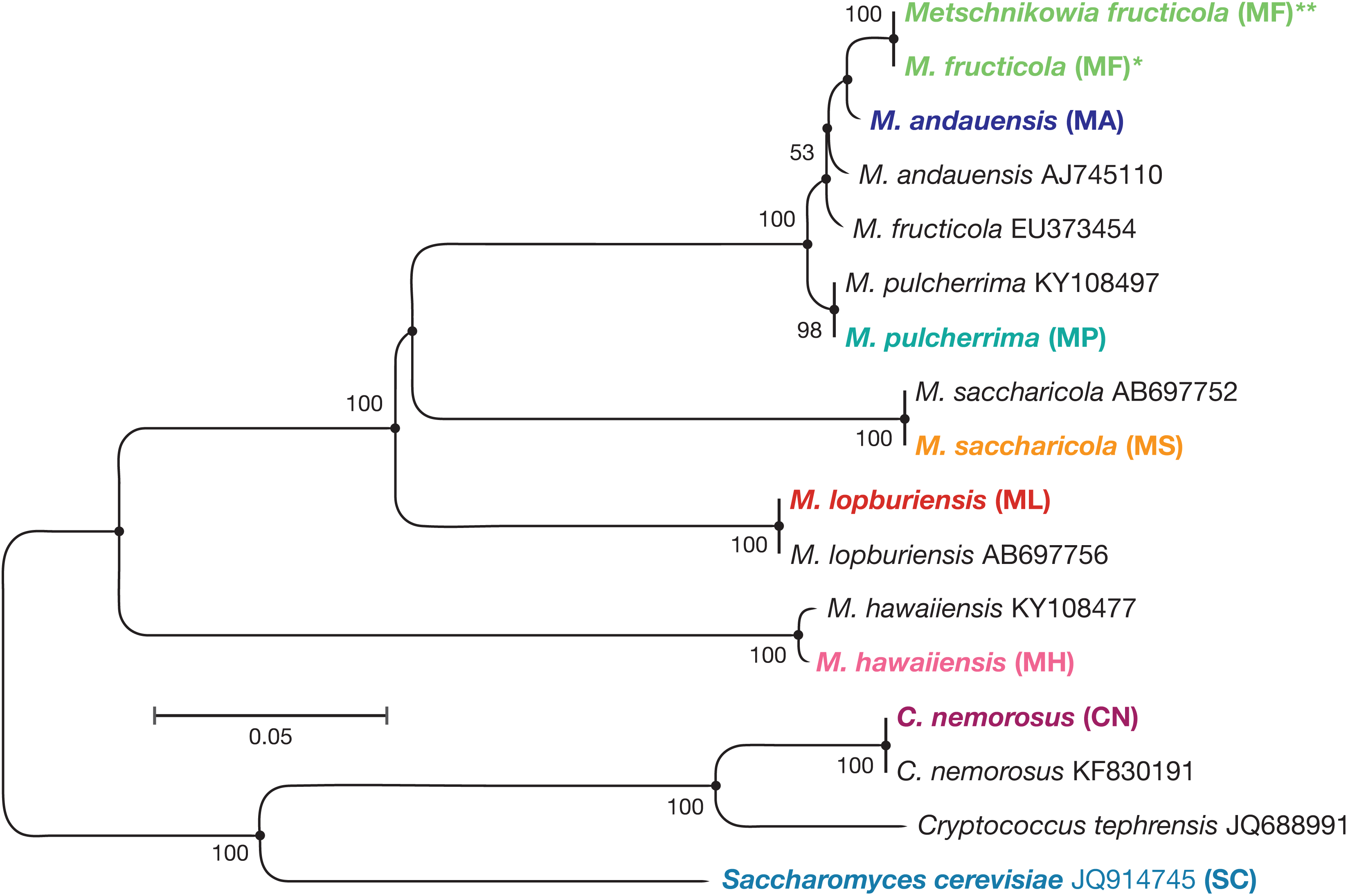
Phylogenetic tree of the yeasts used for volatile analysis (bold-faced) and sequences deposited in the NCBI database, based on the nucleotide sequences of the D1/D2 domain of the 26S rDNA, constructed according to the neighbor-joining method with bootstrap values >50%. Asterisks denote the two isolates of *M. fructicola* corresponding to replicates 1 to 6 and 7 to 12 in Figure 2.

**FIG 2.**
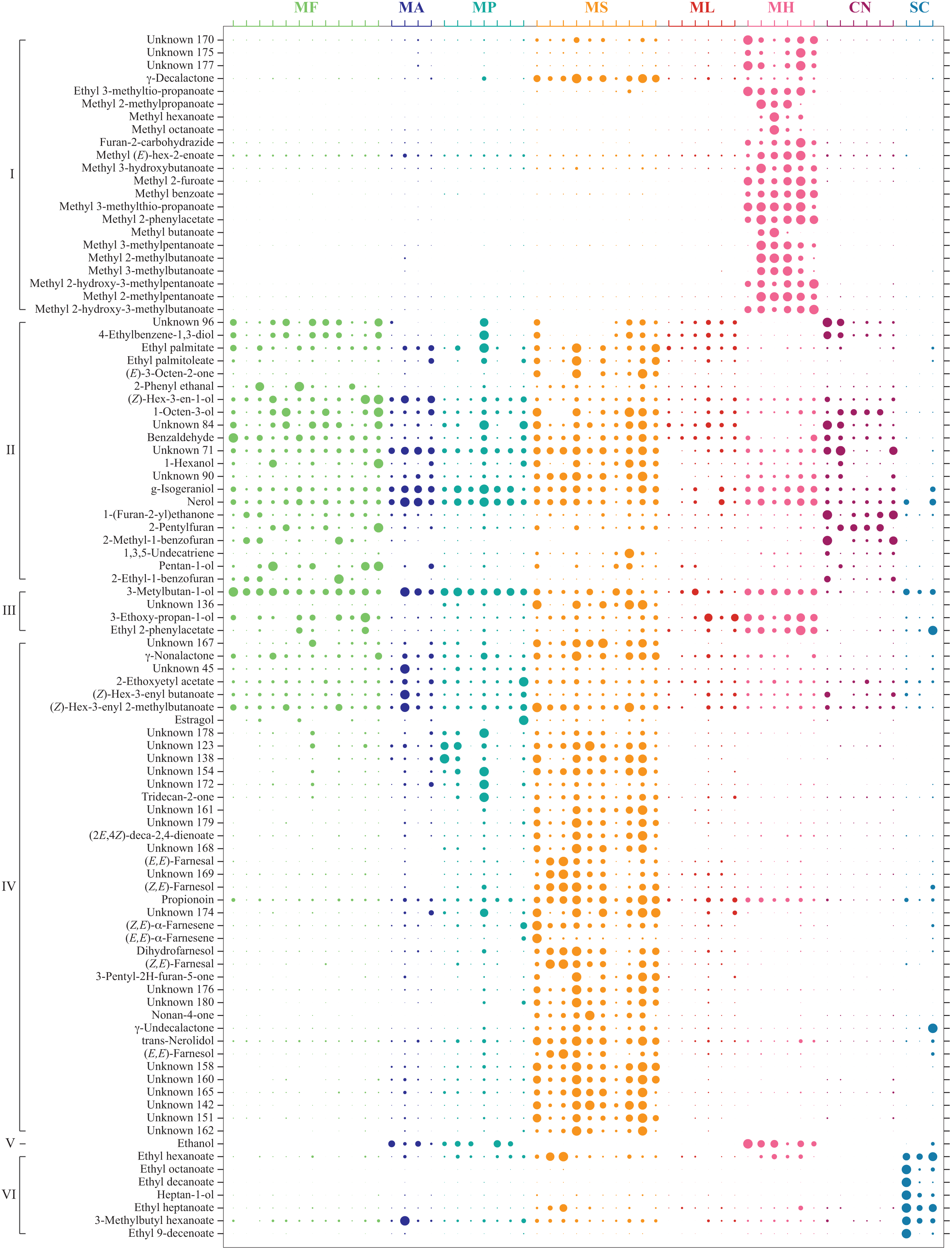
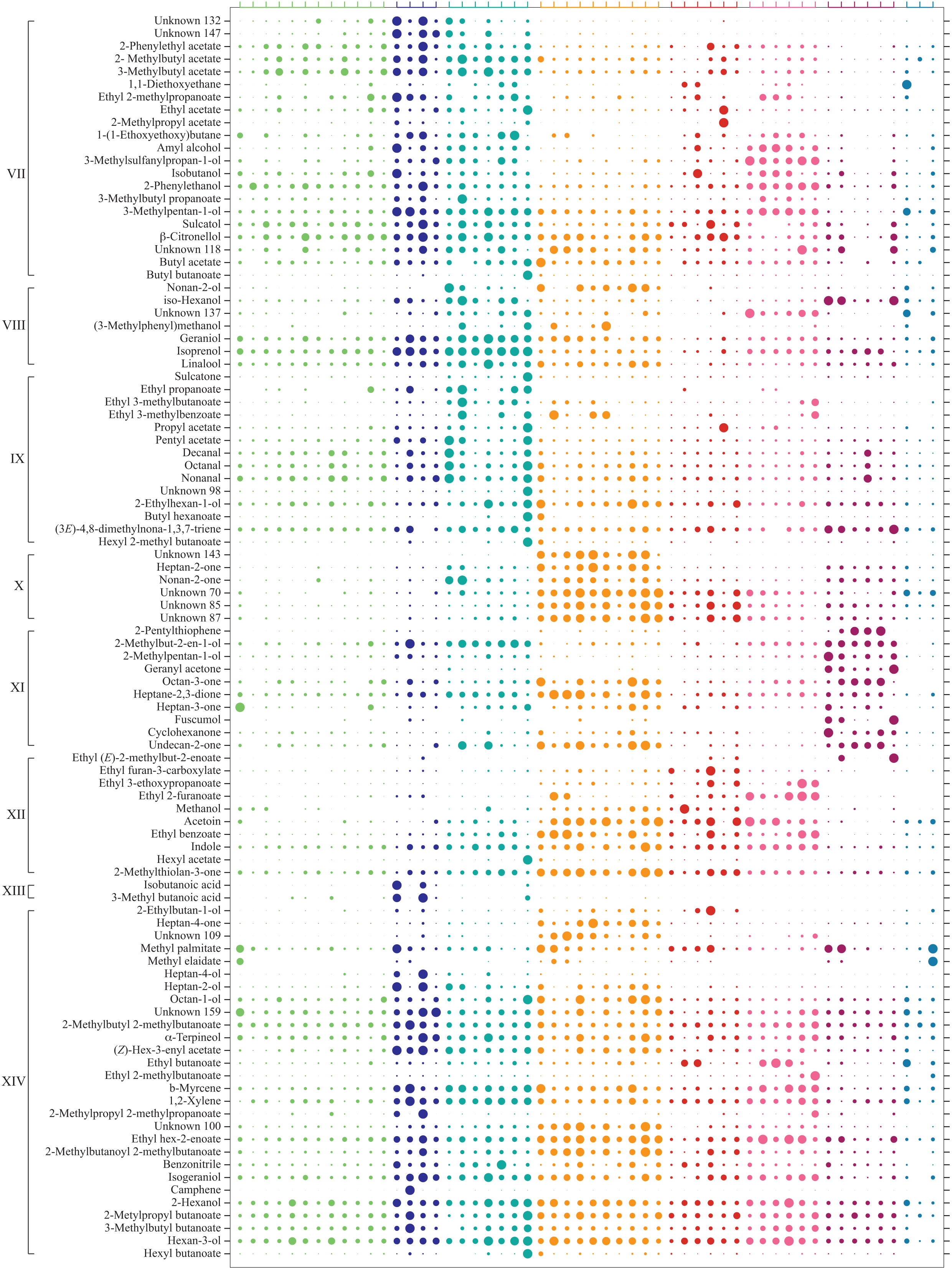
Volatile compounds identified from headspace collections of fermenting yeasts by GC-MS (see also Supplemental Table S1). Compounds were grouped according to hierarchical cluster analysis (HCA), circles depict relative abundance of compounds after normalization by abundance across species (rows) and observations (columns). Yeast species listed according to the phylogenetic tree (Figure 1). Abbreviations see Figure 1.

Figure 2 compares the volatomes of these eight yeasts. Variation across replicates is substantial, despite rigorous protocols employed for yeast growing and headspace collection. However, species could still be separated according to headspace composition by a discriminant analysis (OPLS-DA **M1**, R^2^X_(cum)_ = 0.768; R^2^Y_(cum)_ = 0.918; Q^2^_(cum)_ = 0.756) resulting in 7 predictive components that explain 63% of the entire variation. Especially after grouping compounds into 14 groups by hierarchical cluster analysis (HCA) using the **M1** loadings, Figure 2 further visualises that headspace composition is characteristic for each yeast.

Headspace composition further reflects taxonomic position. *M. andauensis, M. fructicola* and *M. pulcherrima* share morphological and physiological characters, and the D1/D2 domain differs only with respect to few nucleotides (61-63). The close relation between these three species (Figure 1) is in line with their volatome composition, in comparison with the more distantly related *Metschnikowia* species (Figures 2-3).

**FIG 3.**
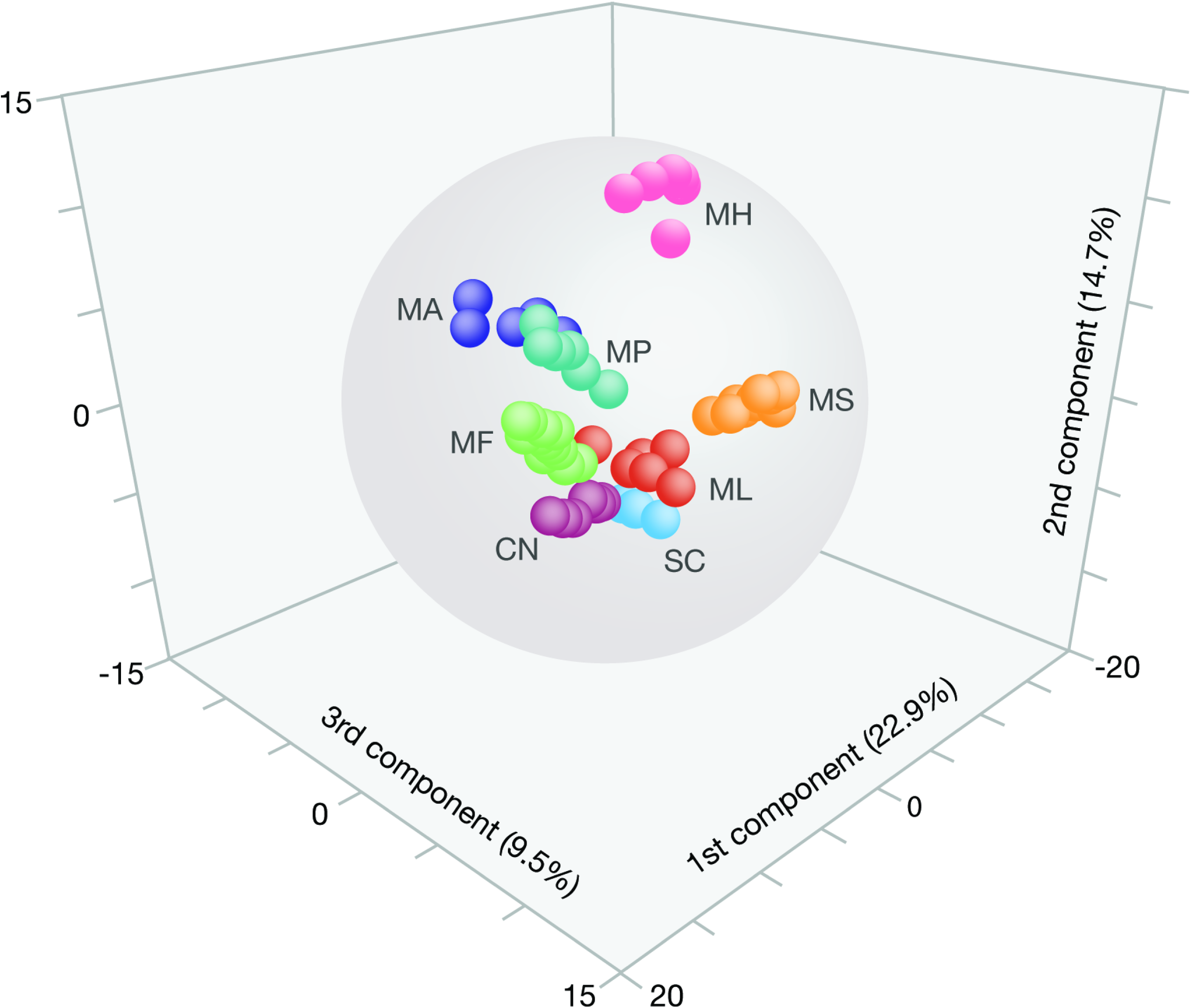
Groups of yeast volatiles according to orthogonal partial least square discriminant analysis (OPLS-DA). The first three principal components represent 22.9 %, 14.7 % and 9.6 % of the total variation in the dataset of eight yeast species. Abbreviations see Figure 1.

Headspace composition also helped to clarify the taxonomic status of a yeast collected from apple, which had been tentatively and incorrectly determined as *Cryptococcus tephrensis*, according to morphological criteria. Visual inspection of its volatome fingerprint suggested this yeast to be closely related to *M. fructicola* (replicates 7 to 12 of *M. fructicola*, Figure 2). This was then confirmed by sequencing the D1/D2 LSU rRNA gene, showing 99% similarity with the sequence obtained from *M. fructicola* (NCBI accession number KC411961, Figure 1).

A three-dimensional score plot of the first three predictive components of **M1** show that *M. hawaiiensis* and *M. saccharicola* clearly separate according to the first three dimensions and that the other species are clustering with little overlap (Figure 3). The remaining species diverged in the remaining dimensions and by HCA of **M1** scores (data not shown). Internal model robustness validation was performed by randomly excluding three observations for each yeast. So as to keep at least three observations, only one replicate was removed for *M. andauensis* and *S.* cerevisiae was removed completely. Excluded observations were thereafter used as prediction set in a new model made of the remaining observations [OPLS-DA **M2**, R^2^X_(cum)_ = 0.686; R^2^Y_(cum)_ = 0.879; Q^2^_(cum)_ = 0.596]. A zero misclassification error (Fishers probability = 2.8 × 10^-10^) further corroborated the robust ability of OPLS-DA to distinguish between yeast headspace profiles.

### *M. hawaiiensis, M. saccharicola* and *M. lopburiensis*

*M. hawaiiensis* has been isolated from morning glory flowers and is associated with drosophilid species (64), and *M. saccharicola* and *M. lopburiensis* have been found on sugarcane and rice leaves (65).

*M. hawaiiensis* and *M. saccharicola* were the most prolific producers of volatiles, several of which were not present in the other yeasts studied (Figure 2). They separated clearly, according to OPLS-DA, from each other and the other yeasts (Figure 3). The headspace of these two species has not yet been studied and contains a number of compounds which are new to databases of yeast volatiles (Supplemental Table S1; 17, 19). They also contained several yet unknown compounds, which were not found in commercial or our own libraries, and which did not match commercially available standards.

Characteristic compounds for *M. saccharicola* were a range of putative sesquiterpenes and also pear ester, a characteristic odorant of pear (66, 67) which is also a strong bisexual attractant for codling moth *Cydia pomonella* (68-70). Indole, a nitrous compound, was released by both *M. lopburiensis* and *M. pulcherrima*. Methanol, ethyl (E)-2-methylbut-2-enoate and ethyl furan-3-carboxylate are the primary class separators for *M. lopburiensis*. Methanol was also consistently found in *M. saccharicola* samples (Figure 2, Supplemental Table S1).

The volatome of *M. hawaiiensis* was clearly separated from the other *Metschnikowia* spp. (Figures 2, 3), methyl esters were key compounds in headspace class separation (Supplemental Table S1). Perhaps coincidentally, two sulphur-containing compounds, methyl 3-methylthio-propanoate and 3-methylsulfanylpropan-1-ol, which are typical for *M. hawaiiensis*, are associated with the aroma of pineapple (71, 72).

### *M. andauensis, M. fructicola* and *M. pulcherrima*

These very closely related species (Figure 1; 63) are morphologically and ecologically similar. They have all been found in larval frass of lepidopteran larvae (41, 62). Three compounds that differentiate *M. fructicola* from other yeasts were 2-phenyl ethanal, 3-ethoxy-propan-1-ol and 3-metylbutan-1-ol. The top four discriminating compounds for *M. pulcherrima* were ethyl 3-methylbutanoate, ethyl propanoate, nonan-2-ol and sulcatone, which showed a high correlation with butyl butanoate. Two methyl-branched short chain carboxylic acids, heptan-4-ol and unknown 147 were highly characteristic for *M. andauensis* (Figure 2, Supplemental Table S1).

### Cryptococcus nemorosus

*Cryptococcus* is polyphyletic and several species such as *C. nemorosus* have been isolated from the plant phyllosphere and soil (73, 74). Ethyl (*E*)-2-methylbut-2-enoate, aliphatic ketones, and aliphatic methyl-branched primary alcohols were the main volatiles that separate *C. nemorosus* from the other species. Strong correlations were observed for 6,10-dimethyl-5,9-undecadien-2-ol (fuscumol) and its respective ketone, 6,10-dimethyl-5,9-undecadien-2-one (geranyl acetone). Shared structures for classifying other species were also observed, namely 2-pentylthiophene was shown to be highly correlating with class membership of *M. andauensis* (Supplemental Table S1).

### Larval feeding on live yeasts

We next investigated attraction and feeding of *S. littoralis* larvae in a choice test. Larval feeding was assayed in a petri dish with two drops of liquid medium, one with live yeast and the other without. Three yeasts, *M. andauensis* (*p* < 0.05), *M. pulcherrima* (*p* = 0.031) and *S. cerevisiae* (*p* = 0.002) deterred feeding, more larvae fed on blank medium in their presence. Two species, *M. fructicola* (*p* = 0.06) and *M. saccharicola* (*p* = 0.096) had no significant effect, but *M. hawaiiensis* (*p* < 0.05), *M. lopburiensis* (*p* = 0.012) and *C. nemorosus* (*p = 0.002*) elicited larval attraction and feeding (Figure 4a).

**FIG 4.**
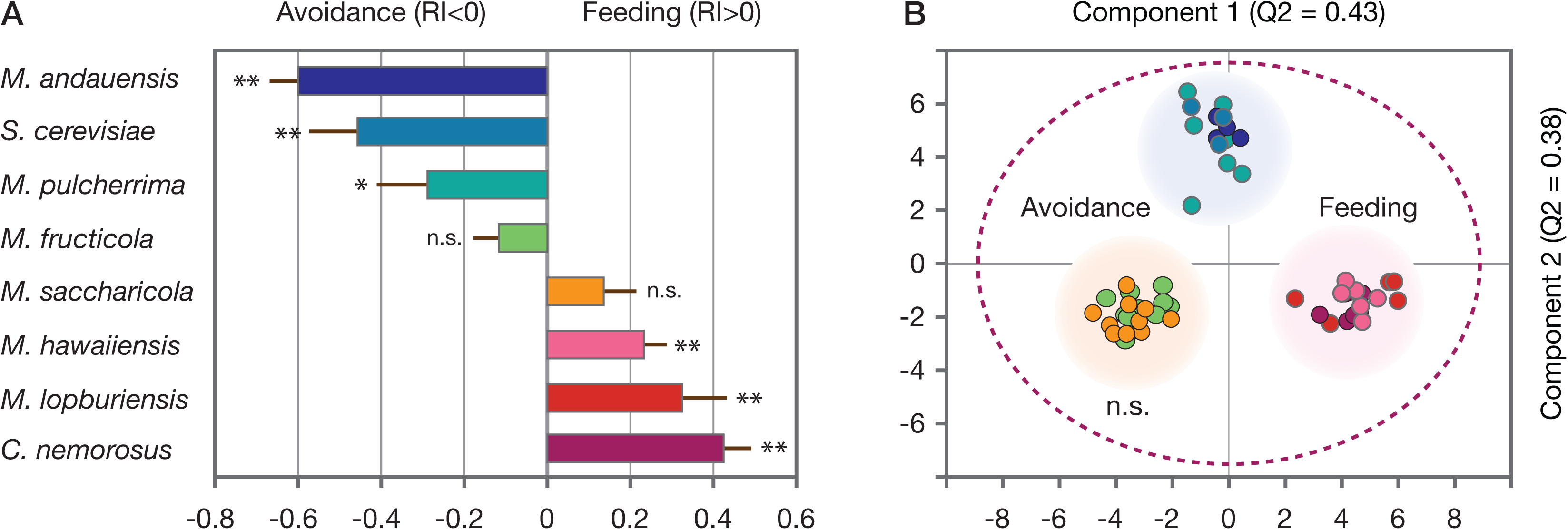
Larval feeding assay and class separation, according to orthogonal partial least square discriminant analysis (OPLS-DA). (a) Bars show the response index RI for larval attraction and feeding (RI>0) and avoidance (RI<0) in response to eight yeasts. Asterisks show significance according to Student’s t-test (P<0.05 and P<0.01, respectively); (b) OPLS-DA score plot of **M3** with yeasts classified according to larvae response. Component 1 and 2 are predictive. Yeasts separate according to behavioral effect, repellency, feeding and no effect. The outline ellipse shows Hotelling’s *T*^2^ (95 %) limit.

An orthogonal partial least squares discriminant analysis (OPLS-DA) was used to explore the correlation of yeast volatiles from different species with respect to behavioural activity. Three classes, shown in Figure 4b, were used which resulted in a model with 2 predictive and 5 orthogonal components [OPLS-DA **M3**, R^2^X_(cum)_ = 0.68; R^2^Y_(cum)_ = 0.947; Q^2^_(cum)_ = 0.812). Model **M3** showed excellent classification performance (Fishers probability = 2.5 × 10^-19^). When plotting the two OPLS-DA predictive components, three groups separate clearly, showing that these yeasts can be distinguished with respect to their behavioural effect (Figure 4b).

Among the eight yeast species, *M. andauensis* and *C. nemorosus* exhibited the strongest activity, resulting in larval avoidance and feeding, respectively (Figure 4a). The volatile profiles of these yeasts show considerable overlap with the other species (Figure 1) and we hypothesized that volatiles released by the other species could be used for sifting inactive volatile constituents and thus facilitate the search for bioactive candidate compounds. We constructed two models to this purpose. Volatiles of all species except *M. andauensis* were modelled against *C. nemorosus*, eliciting the highest rate of feeding which resulted in a model with 2 predictive and 4 orthogonal components [OPLS-DA **M4**, R^2^X_(cum)_ = 0.608; R^2^Y_(cum)_ = 0.984; Q^2^_(cum)_ = 0.933)]. *Metschnikowia andauensis*, which strongly deterred larvae from feeding, was likewise modelled against all other species except *C. nemorosus* [OPLS-DA **M5**, R^2^X_(cum)_ = 0.69; R^2^Y_(cum)_ = 0.983; Q^2^_(cum)_ = 0.572]. Using **M4** and **M5**, a shared and unique structure (SUS) plot was made to illustrate key compounds that are, compared to the other species, released in smaller or larger amounts by *M. andauensis* and *C. nemorosus*. The SUS-plot (Figure 5) assigns two acids, several methyl branched esters, camphene and two unknown compounds 132 and 147 to *M. andauensis*, and geranyl acetone, cyclohexanone, 2-ethyl-1-benzofuran and 1,3,5-undecatriene to *C. nemorosus*.

**FIG 5.**
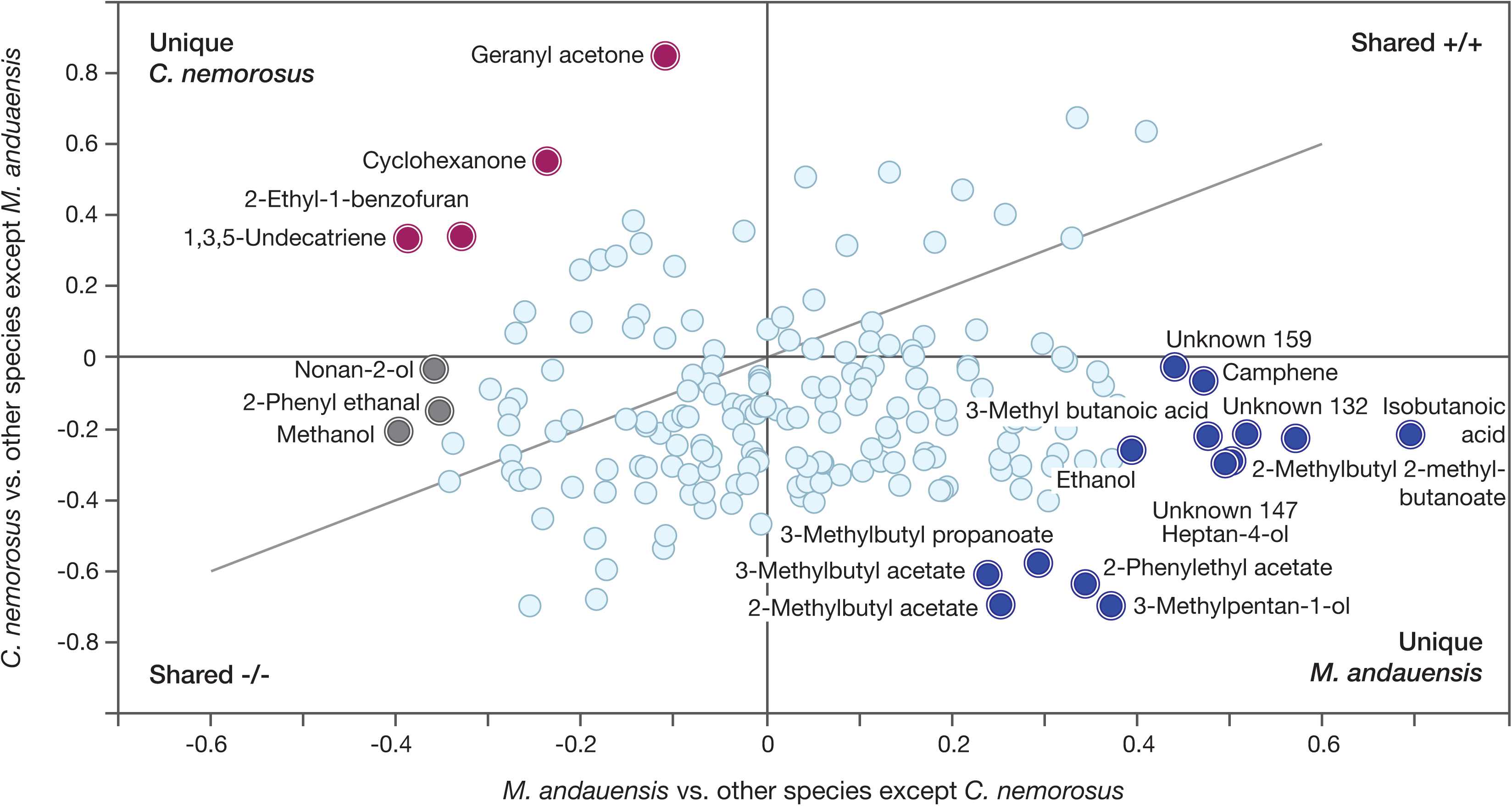
Shared and unique structures plot (SUS-plot) featuring metabolites of the most and least preferred yeasts for cotton leafworm larval feeding, *C. nemorosus* and *M. andauensis* (Figure 4). Correlation from the predictive components of the two models, Corr(*t*_p_,*X*), are plotted against each other. Unique volatiles (top left and bottom right quadrants) are oppositely affected for both *M. andauensis* and *C. nemorosus*. Compounds similarly affected in all yeasts are located along the diagonal running through the shared effect quadrants. Unique, significant volatiles are shown for *M. andauensis* (blue circles) and *C. nemorosus* (red circles).

## DISCUSSION

The yeasts studied here occur on plants, and in connection with insect larvae feeding on these plants. A rich volatome found in all eight species may serve interactions within and between microbial taxa. Presence of many odor-active compounds also supports the idea that yeasts require animal vectors for dispersal and outbreeding. Unlike fungal spores, yeast spores are not adapted for wind-borne transmission. Needle-shaped ascospores, which are frequently found in the *Metschnikowia* clade, promote dispersal by flower-visiting flies, beetles and bees. Yeasts, on the other hand, provide nutritional services to insect larvae and adults (26, 27, 30, 31, 33, 40, 75-78).

Larvae of cotton leafworm *S. littoralis* that naturally feed on foliage of a broad range of annual plants (79, 80) were attracted to volatiles of three yeasts (Figure 4). Figure 2 illustrates that yeast headspace is at least as rich and complex as headspace of their food plants (91, 92), and that also many yeast compounds, including terpenoids, are shared by these plants. This raises the question whether larvae of insect herbivores become attracted to plant or to yeast odour for feeding, or both. Especially for insect species found on a range of plant species, such as *S. littoralis*, volatiles from plant-associated yeasts may be sufficiently reliable signals, especially when these yeasts are part of the larval diet.

The question to which extent yeast vs plant volatiles contribute to oviposition and larval feeding has been formally addressed in *Drosophila melanogaster*, where brewer’s yeast headspace alone elicits attraction and oviposition. *Drosophila* larvae complete their entire development on yeast growing on minimal medium, which supports the conclusion that the fruit merely serves as a substrate for yeast growth (40). In comparison, strict dissection of plant and microbial components is experimentally difficult in insects that require foliage for feeding. For example, grape moth *Paralobesia viteana* was attracted to grape leaves after washing off microbial colonies, but the participating role of microbes remaining on foliage or endophytes is yet unresolved (93).

A closer look at attractants identified from plant hosts produces a surprising insight: typical plant volatiles such as (*Z*)-3-hexenyl acetate, linalool, nonanal or even (3*E*)-4,8-dimethylnona-1,3,7-triene (DMNT), which play an important role in attraction of *P. viteana* or the grape berry moth *Lobesia botrana*, are all produced by several yeasts (Figure 2, 93, 94). Likewise, compounds from cotton headspace that elicit antennal or behavioral responses in cotton leafworm *S. littoralis* are also produced by yeasts (Figure 2, 91, 95). Among these is again DMNT. Induced release of DMNT from plants following herbivore damage attracts natural enemies and deters some insect herbivores. In cotton leafworm, upwind flight to sex pheromone and cotton volatiles is suppressed by large amounts of DMNT, due to its prominent effect on central olfactory circuits (47, 92, 96).

DMNT is also a floral scent component, across a wide range of plants (97) and an attractant of flies and moths (94, 98-100). Yeasts obviously contribute to DMNT release from flowers, since DMNT was found in all yeasts studied here (Figure 2). Besides DMNT, a wide range of volatiles co-occurs in yeasts and angiosperm flowers, for example the typical *Drosophila* attractants acetoin, ethanol, ethyl acetate, 2-phenylethyl acetate, 2-phenylethanol (Figure 2; 101). 2-phenylethanol, a typical yeast odorant (e.g. 102), is also produced by green plants (e.g. 103). Let alone an overlap of compounds produced by both plants and yeasts, fumigation of elderberry flowers with broad-spectrum antibiotics revealed that floral phyllospheric microbiota are unique producers of key floral terpenes (20). The yeasts investigated here all produce a range of terpenes (Figure 2, Supplemental Table S1).

Insect-yeast chemical communication has evolved long before the emergence of flowering plants. Fungivory and herbivory on plants was initiated during the Early Devonian ∼400 Ma, concurrent with the appearance of budding yeasts and prior to angiosperm pollination syndromes during the Cretaceous ∼100 Ma (104-107). This lends support to the idea that a sensory bias for yeast-produced compounds, together with ubiquitous presence of yeasts in flowers, has contributed to the evolution of floral scent and insect-mediated pollination (23, 24, 101, 108, 109).

While the ecological and evolutionary consequences of chemical dialogue between plants, microbes and insects are unequivocal, it is yet largely unclear which of the many volatiles released by yeasts encode this interaction. A comprehensive analysis of yeast volatomes is a first and necessary step towards identifying the active compounds. Thirty-three of the 192 volatiles (Figure 2, Supplemental Table S1) are new for yeasts. Most of them were released by *M. hawaiiensis M. lopburiensis* and *M. saccharicola*, which are the most recently discovered species (64, 65). The database of yeast volatiles creates a basis for future studies, aimed at functional characterization of insect olfactory receptors and attraction bioassays, towards the identification of the behaviourally active compounds (110-112).

The overall species-specific volatome patterns showed variation between replicates, even though growth conditions were strictly controlled and sampling intervals adjusted to cancel out growth stage variations (Figure 2). A general assumption in metabolomics is that identical genotypes produce the same steady state metabolite concentrations under stringent conditions, while metabolic snapshots often show considerable biological variability. Metabolite-metabolite correlations derived from enzymatic reaction network activity may nonetheless be robust, despite considerable intrinsic, stochastic variation of metabolite concentrations obtained at momentary peeks into the state of an organism (113-115)

In spite of inherent variation among volatile samples, numerical headspace analysis by OPLS-DA followed by HCA revealed characteristic volatile fingerprints for each of the eight yeasts (Figures 2, 3) which align with the phylogenetic analysis based on sequences from the D1/D2 region (Figure 1) and yeast taxonomy and ecology (Figure 1; 63, 76). Moreover, we found that OPLS-DA exhibited a robust yeast-species assignment headspace samples and is therefore a useful tool for studying and classifying unknown species with regard to their volatiles (Figure 3).

Volatile fingerprinting or chemotyping has previously been shown to differentiate between ectomycorrhizal, pathogenic and saprophytic fungi, as a complement to genotyping (116, 117). This was confirmed by comparing *M. fructicola* and *M. andauensis*, which are taxonomically close and difficult to discriminate according to genotyping (Figure 1; 118). They quantitatively and clearly separated according to headspace proportions, in addition to production of methanol by *M. fructicola* (Figures 2, 3). Further support for the use of volatomes in species discrimination comes from an isolate from codling moth larvae, which had been misidentified as *C. tephrensis*. Comparison of headspace data with *M. fructicola* evidenced overlap of 101 compounds with significant coefficient values and non-zero confidence interval for the whole dataset. Subsequent DNA analysis identified this yeast as *M. fructicola* (Figure 1).

The species-specific volatome differences are corroborated by a selective larval feeding response (Figure 4), where cotton leafworm larvae, which are typical foliage feeders, prefer phyllosphere yeasts over the yeasts associated with fruit and frugivorous insects. Larvae avoided baker’s yeast commonly found with *D. melanogaster* and *M. andauensis* and *M. pulcherrima* which have been isolated from codling moth, *Cydia pomonella*, feeding in apple (41). It is yet unknown whether cotton leafworm forms yeast associations with yeasts in natural habitats, but a consistent larval response to yeasts may establish and sustain such associations.

Identifying behaviorally active metabolites is key to understanding the ecology of insect-yeast interactions. Geranyl acetone, an aggregation pheromone component of *C. pomonella* larvae (119) is a distinctive compound for *C. nemorosus* (Figures 2, 5), which elicited the strongest larval feeding response (Figure 4). Among the cotton leafworm olfactory receptors which have been functionally characterized, several are tuned to compounds produced by yeasts, and some even elicited larval attraction as single compounds, such as benzyl alcohol, benzaldehyde or indole (120). For a more complete behavioral identification, it would probably be necessary to test compound blends, including candidate compounds from the headspace of *M. hawaiiensis, M. lopburiensis* or *C. nemorosus* (Figures 2, 4, 5).

At the same time, our study reveals potential antifeedants. Camphene has indeed already been reported as a repellant in *S. littoralis* (121) and its acute larval toxicity has been shown in the sister species *S. litura* (122). In addition, *S. littoralis* females detect camphene, as well as 3-methylbutyl acetate (123), both of which are sign compounds for *M. andauensis* (Figures 2, 4, 5). Discriminant analysis points towards presence of attractive and antagonistic yeast volatiles and highlights compounds for future screening assays.

Cotton leafworm is polyphagous on a variety of crops including vegetables in the Afrotropical and western Palearctic. Its sister species *S. litura* is found over Asia, Australasia, and Oceania and the South-American species *S. frugiperda* has recently invaded Africa (52, 60, 79, 124). Global change and increasing food insecurity render insect control an ever more challenging and urgent task (51, 125-127). Detrimental environmental and health effects warrant downregulation of conventional pesticides and accentuate the further development of biological insect control, comprising natural antagonists, insect pathogens or semiochemicals.

Semiochemicals and pathogens are widely and successfully used as stand-alone techniques (128, 129), but semiochemicals could be combined with pathogens into lure-and-kill strategies (57, 58, 130). Current use of insect semiochemicals is based on controlled release formulations of synthetic chemicals, while yeasts could be used for live production of insect attractants (5, 131)

That yeasts would make suitable producers of insect attractants is supported by establishment of biofilms with strong survival ability, which enables postharvest control of fungal diseases in fruit (132-134). Combination of attractant yeasts for targeted ingestion of an insect baculovirus or a biological insecticide has been successful in laboratory and first field experiments, against codling moth and spotted wing *Drosophila* (130, 135, 136). For further improvement, identification of key compounds mediating insect attraction will facilitate selection of yeast species and strains.

Yeast volatiles are also antifungal (132, 133) and may directly, or through other members of the plant microbiome, impact on plant fitness. A critical component of functional interlinkages betwen plants and an ensemble of associated microbiota is that the plant immune system reliably differentiates between synergistic and antagonistic microbes (137, 138). Odorants are essential in regulating mutual and detrimental colonizers in plant microbial networks (6, 7, 139) and yeast volatiles are obviously involved in this chemical dialogue, since compounds such as farnesol or 2-phenylethanol participate in quorum sensing and interspecies interactions (Figure 2, Supplemental Table 1; 16, 140). Integrating plant microbiomes in crop protection concepts, for insect control, enhanced stress tolerance and disease resistance is a future challenge in agriculture (141).

## MATERIALS AND METHODS

### Yeasts

*Metschnikowia* yeasts were purchased from the CBS-KNAW collection (Utrecht, Netherlands), except *M. fructicola*, which was isolated from apple (Alnarp, Sweden) infested with larvae of codling moth *Cydia pomonella* (Lepidoptera, Tortricidae), *Cryptococcus nemorosus* was isolated from cotton leafworm *Spodoptera littoralis* (Lepidoptera, Noctuidae) larvae (laboratory rearing, Alnarp, Sweden), and *Saccharomyces cerevisiae* was obtained from Jästbolaget AB (Sollentuna, Sweden).

*M. andauensis, M. fructicola* and *M. pulcherrima* were found in guts and larval feces of caterpillars feeding on maize, corn earworm *Helicoverpa armigera* (Lepidoptera, Noctuidae) and European corn borer *Ostrinia nubilalis* (Lepidoptera, Crambidae) (62). *M. andauensis* and *M. pulcherrima*, and occasionally *M. fructicola* were found in apple and larval feces of codling moth *Cydia pomonella* (Lepidoptera, Tortricidae) (41). *M. saccharicola, M. lopburiensis* were isolated from foliage in Thailand (65) and *M. hawaiiensis* from fruit flies (Drosophilidae, Diptera) (64).

### DNA isolation and yeast identification

Genomic DNA was isolated from overnight yeast cultures grown in liquid yeast extract peptone dextrose (YPD) medium. 2 mL of the overnight cell cultures were pelleted (13.000 rpm for 1 min) and washed with sterile ddH_2_O. The pellets were resuspended in 200 μL lysis buffer (2% Triton X-100, 1% SDS, 0.1 M NaCl, 10 mM Tris, 1 mM EDTA, pH 8), and 200 μL phenol:chloroform:isoamyl alcohol (25:24:1) mixture. The glass beads (200 µL) were then added to the tubes and vortexed thoroughly for 3 min. To this mixture, 200 μL TE buffer (10 mM Tris-Cl, 1 mM EDTA; pH 8.0) was added and centrifuged at 13.000 rpm for 10 min. The upper aqueous phase was collected and 10 μL of RNaseA (10 mg/mL) was added and incubated at 37°C during 45 min. The DNA was precipitated using 300 mM sodium acetate, three volumes of cold absolute ethanol (99.9%) and centrifuged at 13.000 rpm for 10 min at 4°C. The DNA pellet was then washed in 70% cold ethanol (13.000 rpm for 10 min, 4°C), air-dried, and re-suspended in 30 μL TE buffer and stored at −20°C until use.

The D1/D2 domains of the 26S ribosomal RNA gene was amplified with the universal primer pair NL-1 and NL-4 and the internally transcribed spacer (ITS) region with the primer combination ITS-1 and ITS-4 (142, 143). The PCR reaction and product visualisation on agarose gel was performed as previously described (101). PCR-products were purified using ExoStar-IT (USB Corporation, USA) following the manufacturer’s protocol sequenced using BigDye v.1.1 terminator sequencing kit in an ABI PRISM 3130 genetic analyser (Applied Biosystems, New Jersey, US).

All sequences were aligned in the program Bioedit 7.0.9.0 Sequence Alignment Editor (144). Each sequence was tested for identity and similarity against sequences deposited in the National Center for Biotechnology Information (NCBI) using Megablast-search (accession number KF839191, KC411961). Only similarities over 95% were considered. The phylogenetic tree of the yeast species investigated, based on nucleotide sequences of the D1/D2 domain of 26S rDNA, was constructed using the neighbor-joining (NJ) method in MEGA 7.0 (145). The NJ method, based on the evolutionary distance data that minimizes the total branch length during clustering of operational taxonomic units (OTUs), is efficient and reliable for phylogenetic reconstructions (146). The evolutionary distances were calculated according to the Jukes and Cantor (JC) substitution model to minimize the bias due to nucleotide substitution during divergence. The JC substitution model considers the rate of substitution frequencies of all pairs of four nucleotides (A, T, G, C) to be equal (147). The bootstrap values for the phylogenetic tree reconstruction were determined from 1.000 replications and are given next to the branch (Figure 1).

### Headspace collection and chemical analysis

Yeasts were grown in 100 mL liquid minimal medium (148) in 250-mL culture flasks, during 24 h in a shaking incubator (25°C, 260 rpm). Yeast headspace was collected by drawing charcoal-filtered air (0.125 L/min), through a 1-L gas wash bottle containing the yeast broth, over a 35-mg Super Q trap (80/100 mesh; Alltech, Deerfield, IL, USA), which was held between plugs of glass-wool in a 4 × 40-mm glass tube. Collections were done for ca. 24 h, at 20 to 22 °C and 10 to 30 Lux. The charcoal filter (50 g activated charcoal) for incoming air and the Super Q trap were connected with glass fittings to the wash bottle. All glassware was heated to 375°C for 10 h before use (149).

Following volatile collections, the trap was extracted with 0.5 mL of redistilled hexane. Sample volumes were reduced to ca. 50 µL, at ambient temperature in Francke-vials with an elongated tip (5 cm × 2 mm i.d.). Samples were stored in sealed glass capillary tubes at −19°C. The Super Q trap was wrapped in aluminium foil for protection from light. Before use, it was rinsed sequentially with 3 mL of methanol (redistilled >99.9% purity, Merck, Darmstadt, Germany) and hexane (redistilled >99.9% purity; Labscan, Malmö, Sweden).

Yeast headspace collections were analysed on a coupled gas chromatograph–mass spectrometer (GC–MS; 6890 GC and 5975 MS; Agilent Technologies, Palo Alto, CA, USA), operated in the electron impact ionization mode at 70 eV. The GC was equipped with fused silica capillary columns (30 m x 0.25 mm, d.f. = 0.25 µm), DB-Wax (J&W Scientific, Folsom, CA, USA) or HP-5MS (Agilent Technologies). Helium was used as the mobile phase at an average linear flow rate of 35 cm/s. Two µL of each sample were injected (splitless mode, 30 s, injector temperature 225°C). The GC oven temperature for both columns was programmed from 30°C (3 min hold) at 8°C/min to 225°C (5 min hold).

Data were exported in NetCDF file format and deconvoluted into compound spectra, elution profile and peak area. We used MS-Omics software (Vedbaek, Denmark) and the PARAFAC2 model (150), and the non-commercial package HDA (version 0.910, P. Johansson, Umeå University, Sweden), based on the H-MCR method (151). Both methods utilize covariation between samples to separate co-eluting components, and to pool mass spectra across samples affording unambiguous spectra even at low signal to noise ratios. Approx. 70% and 30% of the peaks were separated by PARAFAC2 and H-MCR, respectively. However, neither deconvolution method produced satisfactory results for all compounds, which necessitated manual selection of the most feasible method in some cases.

Compounds were identified according to their retention times (Kovat’s indices) and mass spectra, in comparison with the National Institute of Standards and Technology (NIST, version 14) mass spectral library and authentic standards, on two columns. Extra care was taken to verify the identity of compounds showing high variation in abundance between yeast species. Compounds present in blank recordings of the growth medium were subtracted.

### Insects

A cotton leafworm *Spodoptera littoralis* (Lepidoptera, Noctuidae) laboratory colony was established using field-collected insects from Alexandria (Egypt) in 2010. This colony was interbred with wild insects from Egypt every year. Insects were raised on a semisynthetic agar-based diet (152) under a 16L:8D photoperiod, at 24°C and 50 to 60% RH.

### Larval feeding assay

Yeasts (*M. andauensis, M. fructicola, M. hawaiensis, M. lopburiensis, M. pulcherrima, M. saccharicola, S. cerevisiae* and *C. nemorosus*) were grown in 125-mL culture flasks in 50 mL of liquid minimal medium (148) for 20 h in a shaking incubator (25 °C, 260 rpm). Optical density at 595 nm was between 1.5 and 1.8, and cell counts were adjusted to 1.5 × 10^7^ cells/mL.

A two-choice bioassay was conducted to determine neonate larval yeast attraction and feeding. Two 50 µL drops of a 20-h old yeast culture and blank minimal medium were pipetted opposite from each other, approx 1 cm from the edge of a plastic petri dish (92 mm ø, No.82.1472, Sarstedt AG & Co., Nümbrecht, DE). Colorants, blue and green, (Dr. Oetker Sverige AB, Göteborg, SE) were used at 1:10 dilution to color yeast and minimal medium, in order to distinguish between the larvae that fed on yeast medium, blank medium or both. Preliminary tests did not show a bias in larval attraction to the different colorants (N=30; F=1.529; p=0.222; ANOVA).

Ten starved neonate larvae were collected 24 to 36 h after hatching and were placed in the centre of the petri dish with a fine brush. The dish was covered with a lid to prevent the larvae from escaping, and larvae were left to feed for 2 h. They were then checked under a microscope for their gut coloration. Ten independent replicates with 10 neonate larvae were done for each yeast. The larvae response index (LRI) was calculated from the number of larvae feeding on the yeast treatment (nY) and control (nC): LRI = (nY-nC)/(nY+nC).

### Statistical Analysis

All yeast species and volatile compounds were collated into a matrix, containing 56 yeast headspace collections and the integrated areas of the 192 compounds found. Two pre-treatment methods were selected before building the model. A logarithmic transformation of x-variables served to minimize skewness of the variables. Since the abundance of compounds does not necessarily correlate with biological activity, pareto scaling was used to augment the impact of compounds present in very small amounts. Hierarchical cluster analysis (HCA), orthogonal partial least square discriminant analysis (OPLS-DA) and partial least square discriminant analysis (PLS-DA) and its corresponding validation tests were calculated with SIMCA 13.0.3 (Sartorius Stedim Data Analytics AB, Umea, Sweden).

Three methods were used to validate the multivariate models. Internal cross validation (CV) was done with eight CV-groups of dissimilar observations. A permutation test, by randomization of the class vector, was used to test for the presence of spurious correlations between volatile profiles and class membership by fitting each permutation to a new PLS-DA model and observing the resulting explained variance and predictive power. Finally, an internal validation set of observations that spanned the multivariate space for each model was inspected for any large deviations.

Larval feeding of *S. littoralis* on yeasts was compared using Student’s t-test, with the level of significance set to p=0.05.

## SUPPLEMENTAL MATERIAL

Supplemental material for this article may be found at …

**SUPPLEMENTAL TABLE S1**, PDF file, 0.15 MB.

## ACKNOWLEDGEMENTS

This work was supported by the Linnaeus environment “Insect Chemical Ecology, Ethology, and Evolution (IC-E3)” (Formas, SLU), Västernorrland County Administrative Board, European Regional Development Fund of the European Union and the Faculty of Landscape Architecture, Horticulture and Crop Production Science (SLU, Alnarp). The authors gratefully acknowledge the Swedish Metabolomics Centre, Umeå for access to GC-MS evaluation software and helpful discussions.

---

**TABLE S1.**
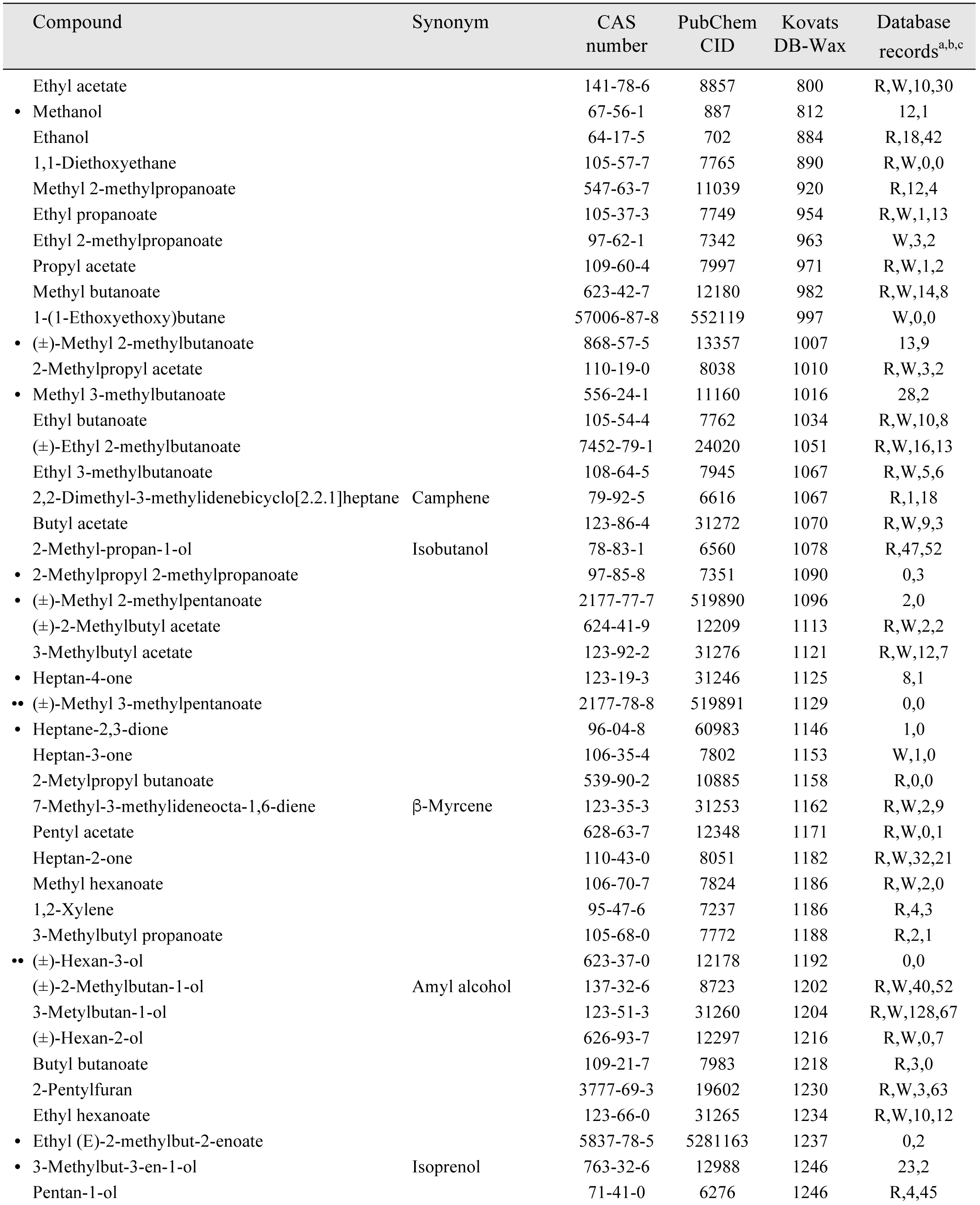
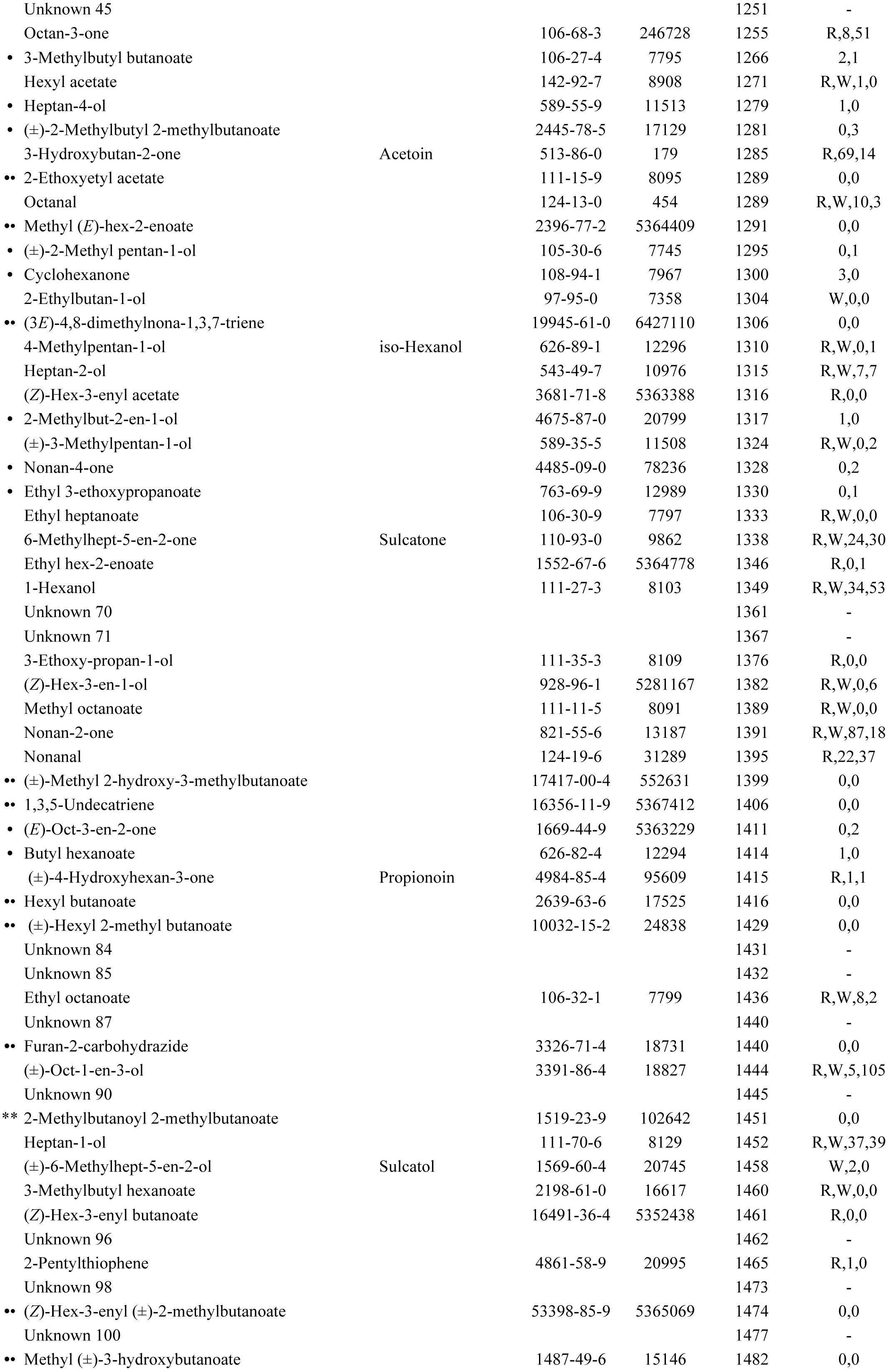
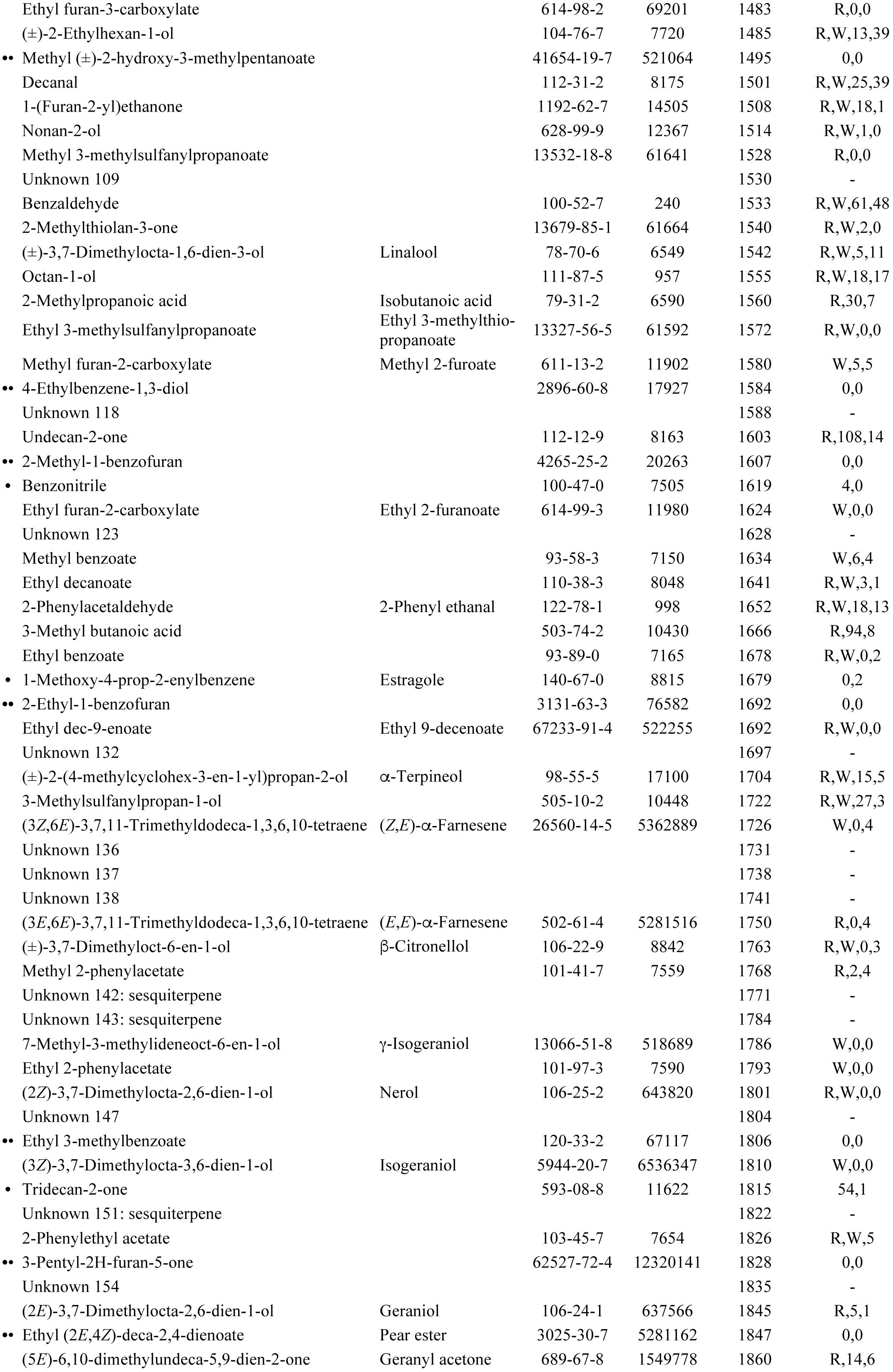
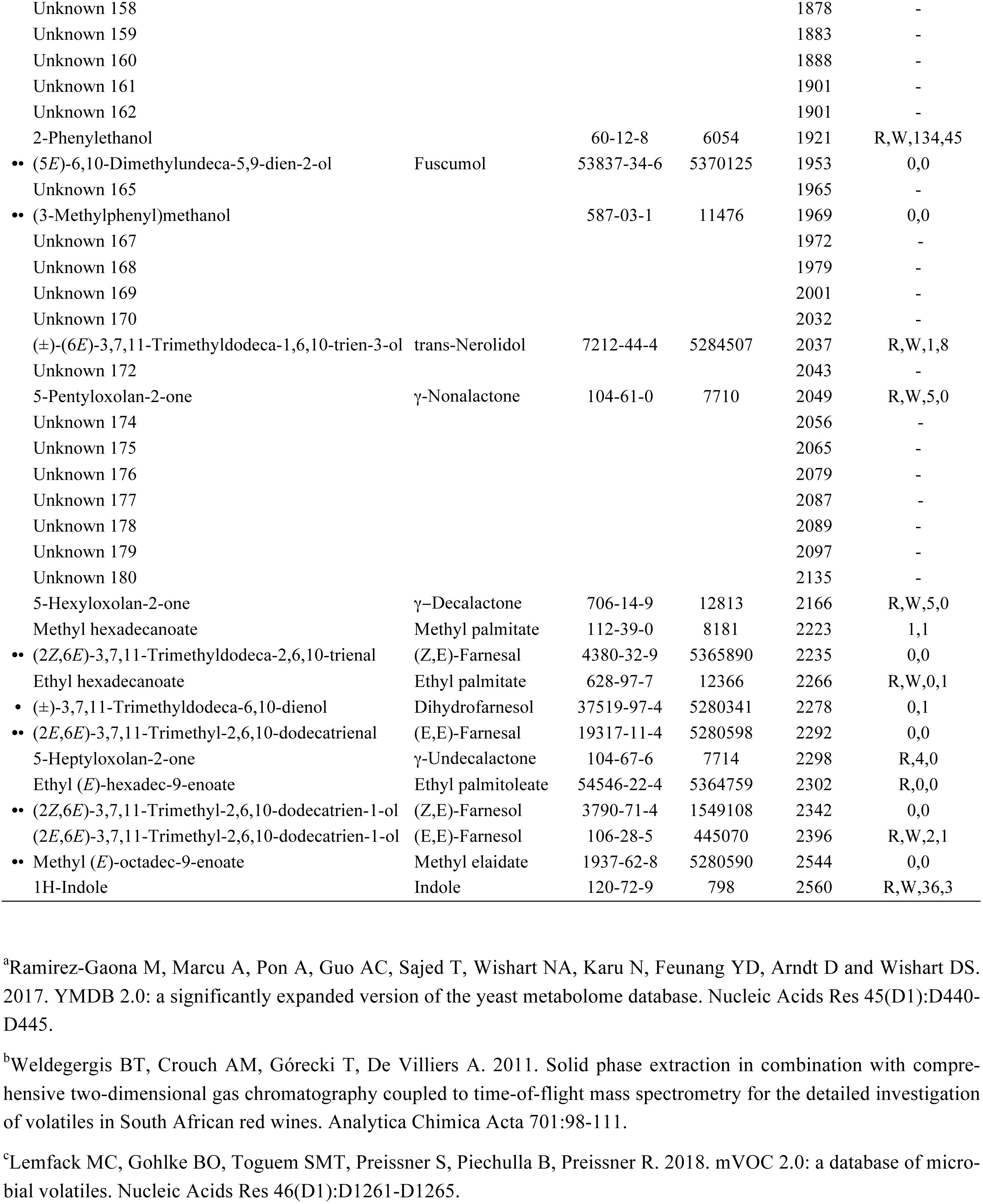
Volatile compounds identified by GC-MS from the headspace of eight yeasts, *Cryptococcus nemorosus, M. andauensis, M. fructicola, M. hawaiiensis, M. lopburiensis, M. pulcherrima, M. saccharicola* and *Saccharomyces cerevisiae*. Compounds marked with a bullet are not listed in yeast databases, two bullets for compounds not known from databases of yeast, bacterial or fungal metabolites. Compound nomenclature follows IUPAC, according to PubChem. CAS and CID number are shown, and Kovats indices on a DB-Wax column. Letters R and W denote compounds listed in yeast metabolome databases by Ramirez-Gaona et al. (2017)_a_ and Weldegergis et al. (2011)_b_, followed by the number of bacteria and fungi, respectively, producing these compounds according to Lemfack et al. (2018)_c_.

